# Imaging Ultraweak Photon Emission from Living and Dead Mice and from Plants under Stress

**DOI:** 10.1101/2024.11.08.622743

**Authors:** V. Salari, V. Seshan, L. Frankle, D. England, C. Simon, D. Oblak

## Abstract

The phenomenon of biological ultraweak photon emission (UPE), that is, extremely low-intensity emission (10 *−* 10^3^ photons/cm^2^/sec) in the spectral range of 200 *−* 1000 nm, has been observed in all living systems that have been examined. Here we report experiments that exemplify the ability of novel imaging systems to detect variations in UPE for a set of physiologically important scenarios. We use EMCCD and CCD cameras to capture single visible-wavelength photons with low noise and quantum efficiencies higher than 90%. Our investigation reveals significant contrast between the UPE from live vs. dead mice. In plants we observed that an increase in temperature and injuries both caused an increase in UPE intensity. Moreover, chemical treatments modified the UPE emission characteristics of plants, particularly the application of an anesthetic (benzocaine) to injury, which showed the highest emission among the compounds tested. As a result, UPE imaging provides the possibility of non-invasive label-free imaging of vitality in animals and the responses of plants to stress.

## I. INTRODUCTION

Ultraweak photon emission (UPE), also known as biophoton emission, refers to the spontaneous emission of extremely low levels of light from all biological systems such as cell cultures, bacteria, plants, animals, humans, etc [1–11]. It was initially observed in plants and later in animals and human cells. However, following decades of research, the origins and significance of this emission is still not completely understood. It is known that reactive oxygen species (ROS) play an important role in the phenomenon of UPE [12]. When organisms face stressors such as environmental toxins, pathogens, or physiological imbalances, they often trigger biochemical pathways that result in the production of ROS, such as, superoxide anion, hydrogen peroxide, and hydroxyl radical, which are signaling molecules in cellular responses to stress [13]. Excessive ROS production can overwhelm the cellular antioxidant defense mechanisms, leading to oxidative stress [14– 18]. ROS-induced lipid peroxidation and protein oxidation can trigger electron excitation and transfer, leading to UPE [19–21]. Moreover, ROS-mediated reactions can alter the redox state of cellular components, influencing the dynamics of radical pair reactions and subsequent photon emission [22– 28]. The correlation between stress in biological systems and the generation of ROS is a key aspect of cellular physiology [13]. For instance, the ability to monitor UPE could potentially enable early detection of stress-related pathologies, e.g. cancer [45, 46, 68–70].

The endeavor to detect single photons began with the help of Photomultiplier Tubes (PMTs), which were pioneering systems but with limited sensitivity and spatial resolution[29]. Technological advancements in equipment, such as, Charge-Coupled Devices (CCDs) [30–32, 34, 35] and Electron Multiplying CCDs (EMCCDs) cameras[36–38] for imaging, significantly boosted the ability to capture single photons with higher efficiency and resolution. Using these tools, UPE imaging has been extensively investigated across various living systems [44]. For instance, significant contributions have been made using CCD cameras to image UPE from different living systems [45–48], e.g. human UPE has been studied for physiological and pathological states [49], and UPE imaging has been performed to investigate the links with reactive oxygen species [22, 23, 50]. Additional imaging studies span multiple systems with a particular interest in the biophysical mechanisms underlying UPE [51–53].

To the best of our knowledge, there have been no reports of the impact of mortality on UPE through imaging thus far. Additionally, while the scientific literature has touched upon the influence of temperature on variations in UPE [54], this effect has rarely been imaged. Moreover, although a few studies have explored the impact of stress [43] and injury [71] in plants, the effects of chemicals such as anesthetics on injury have not been previously investigated. In this paper, we present the results of our experimental investigation on living and dead mice to examine the mortality effect on UPE and elucidate the significance of UPE under different stress effects such as temperature, injury, and chemical treatment in plants.

## II. SPONTANEOUS BIOLOGICAL LIGHT EMISSION

Spontaneous emission in biological systems is the phenomenon where organisms emit light without requiring external excitation. After being in darkness for a period, these organisms still produce light. This type of emission occurs naturally and is typically a result of specific chemical reactions within the organism’s cells or temperature. There are three types of spontaneous light emission from living systems: blackbody radiation, bioluminescence, and UPE [66].

Blackbody radiation (BBR), or thermal radiation, refers to the electromagnetic radiation emitted by any surface due to its temperature in thermal equilibrium, and has an intensity that increases with temperature. Bioluminescence is high-intensity light emission (10^5^ and higher orders photons/cm^2^ per sec) by living organisms that can be seen by the naked eye. It typically involves the enzyme luciferase and a substrate such as luciferin, or can be due to non-enzymatic reactions such as photoproteins, found in certain organisms, e.g. bacteria, fireflies, and jellyfish, and serves various purposes including communication, camouflage, and attracting prey or mates. The UPE is the lowest intensity light that can be observed from all living systems with only the most sensitive detectors. It typically occurs within the near-infrared, visible, and ultraviolet range of the electromagnetic spectrum and is believed to originate from various biochemical reactions and metabolic processes within cells [11, 56]. As we mentioned earlier, UPE can serve as a sensitive indicator of cellular metabolism, oxidative stress, and physiological well-being [12].

UPE and blackbody radiation are present for all organisms, but have distinct characteristics. Blackbody radiation follows Planck’s law [57] and depends solely on temperature, with a wavelength spectrum mostly ranging from wavelengths significantly above 1000 nm at room temperature. UPE, on the other hand, originates specifically from biological systems and falls in spectral ranges determined by specific biochemical processes, reportedly mostly below 1000 nm. While UPE can be affected by temperature variations [54] it can also be modulated by various other factors, including changes in cellular activity, biochemical reactions, and the presence of certain chemicals or molecules within living organisms. This dynamic nature of biological processes and photon emission sets UPE apart from the more static phenomenon of blackbody radiation. We can obtain a lower bound of the spectral energy density of the observed radiation by numerically integrating the energy of the photon flux over a spectral range of 200 nm to 2000 nm for blackbody radiation and 200 nm to 1000 nm for ultraweak photon emission (UPE). For comparison, the spectral energy density of black bodies at ambient (22°C) and body (37°C) temperature can be calculated by Planck’s law [57]:

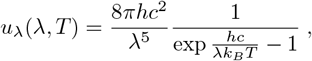

where *u*_*λ*_(*λ, T*) is the spectral energy density from a black body at a specific wavelength *λ* and temperature *T*. Fig.1 shows that UPE is orders of magnitude higher than the black body radiation at temperatures typically present in UPE imaging setups. As a result, the effect of black body radiation is negligible compared to UPE in the spectral range observable by our imaging sensors.

**FIG. 1.**
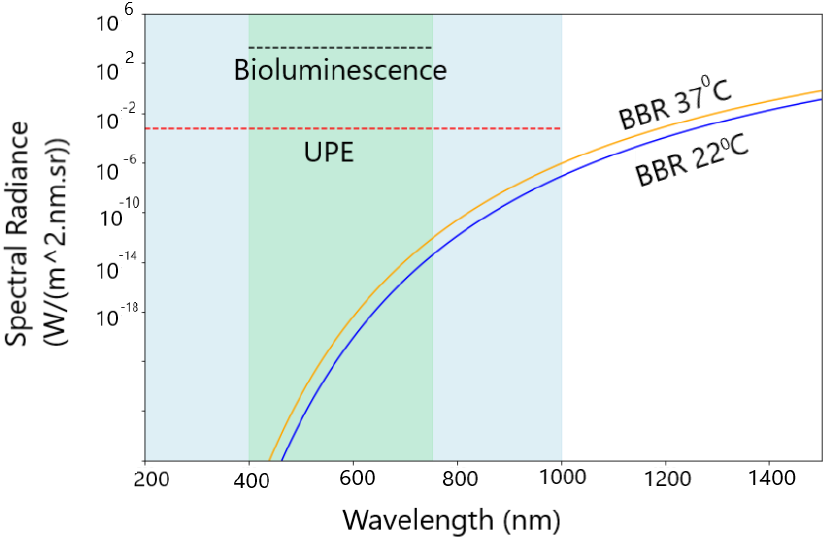
Photon fluxes of blackbody radiation (BBR) for UV-VisNIR wavelengths at room temperature (T=22°C) and body temperature (T=37°C) along with the minimum spectral energy density of UPE (assuming a minimum photon flux of 10 photons/(cm^*−*2^ s^*−*1^)) and the estimated spectral energy density of bioluminescence (assuming a minimum photon flux of 10^5^ photons/(cm^*−*2^ s^*−*1^)) which is limited to the visible spectrum [10, 11, 66].

## III. EXPERIMENTAL METHODS

High quantum efficiency CCD and EMCCD cameras are among the most sensitive single photon detection devices, particularly for UPE imaging and studies [34, 35, 38]. A CCD camera is a type of image sensor commonly used in digital cameras and imaging devices that captures photons and converts them into electrical signals, creating digital images. The high quantum efficiency CCD cameras are used for scientific research [30–32]. In contrast, an EMCCD camera is a specialized type of CCD camera that includes an electron multiplication register, allowing for extremely sensitive detection of low-light signals. This makes EMCCD cameras ideal for applications requiring high sensitivity and low-light imaging, such as astronomy and bioluminescence imaging [36, 37].

The EMCCD technology amplifies weak signals while maintaining high quantum efficiency.

### 1. Cameras

In this research, we have used two cameras for imaging, one EMCCD camera Andor iXon ultra 888 at the Quantum Cloud Lab at the University of Calgary for imaging plants under stress, and a CCD camera at the Human Health Therapeutics Research Center, National Research Council Canada (NRC), Ottawa, which is an IVIS imaging system equipped with a CCD camera Andor iKon L series (see Fig.2) for imaging mice. The EMCCD iXon 888, and CCD iKon L series boast a remarkable quantum efficiency exceeding 95% in the visible range (see Fig.2) with low dark current rates 0.00011 and 0.000059 (e-/pixel/sec) at max cooling respectively.

**FIG. 2.**
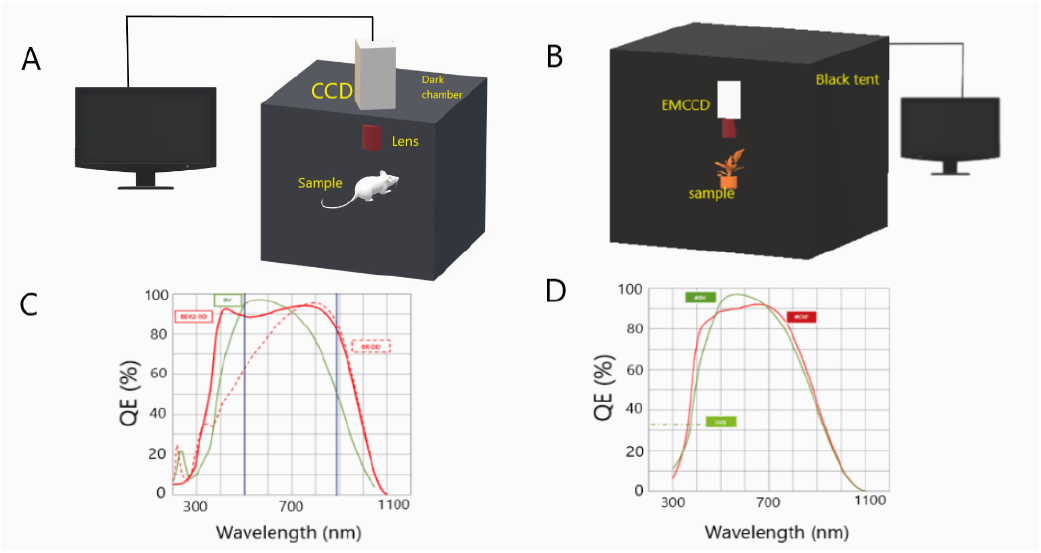
The imaging setups at the NRC (A) based on an IVIS Lumina imaging system equipped with a CCD camera, Andor iKon L-series, with quantum efficiency diagram (C) for the standard Silicon (‘BV’) sensor, and the setup at the University of Calgary (B) featuring a large dark-tent housing the imaging setup equipped with an EMCCD camera, Andor iXon ultra 888, with quantum efficiency diagram for the BV sensor (D). The BV sensor features a high quantum efficiency in the visible range. The QE diagrams are taken from the manufacturer’s specifications of the cameras[33]. The experiments on mice were performed at the NRC and the experiments on plants were done at the UCalgary.

The EMCCD camera was connected to a computer via a USB cable and data acquisition was conducted with the help of the Solis software. For enhanced sensitivity and reduced dark current, an external water chiller was connected to the sensor to achieve a stable operating temperature of -95°C. The images were taken with vertical shift speed *µ*s, electron-multiplying gain mode 2 and amplification gains 300 and 1000 for different experiments. To improve the signal-to-noise ratio, 2 *×*2 and 4 *×*4 binning were used along with a cosmic ray filter. The cameras were placed in ultra-dark enclosures in dark rooms to remove environmental light penetration (see Fig.2).

#### A. Samples

##### 1. Mice

UPE imaging on mice was performed using an IVIS Lumina III preclinical animal imager (PerkinElmer, Waltham, Massachusetts, USA). SKH1 Elite hairless mice (Charles River, Montreal, Quebec, Canada) were anesthetized with isoflurane gas and positioned on the imaging stage, where anesthesia was maintained for the duration of the imaging session. Before imaging, mice underwent a 30-minute dark acclimation period, followed by a 60-minute biophoton imaging session in the dorsal position with the following parameters: single mouse position and open filter. After the imaging session, mice were euthanized using CO_2_. After euthanasia, the mice were immediately placed in the imager, acclimated to darkness for 30 min and then re-imaged using the same settings. Total photon flux was quantified using Living Image 4.1 software (PerkinElmer, Waltham, Massachusetts, USA), see Fig.2.

##### 2. Plants

For UPE imaging of plants and leaves, live Arabidopsis thaliana plants and freshly cut green leaves from Heptapleurum arboricola plants at the UCalgary were used. We investigated the effects of injury and different chemical agents on the injury response of plants. The chemicals used were alcohol (isopropanol), hydroge peroxide (H_2_O_2_), and benzocaine. Alcohol is a solvent and disinfectant that can permeabilize cell membranes and denature proteins. When applied to injured plant tissues, alcohol may induce stress responses by compromising cellular integrity and triggering signaling pathways involved in defense and repair mechanisms. This can lead to alterations in reactive oxygen species (ROS) levels, impact metabolic processes, and affect intercellular communication, depending on the concentration and exposure duration. Benzocaine is a local anesthetic known to inhibit nerve impulses by blocking voltage-gated sodium channels. In plants, benzocaine can potentially disrupt ion channel activities [39] or interfere with signaling pathways involved in stress responses or with a different mechanism related to ROS[40–42]. Hydrogen peroxide is a type of ROS whose amount has a direct correlation with UPE intensity. It is also a key signaling molecule involved in plant responses to stress and injury [43]. It acts as a secondary messenger, to trigger various defence and repair pathways.

Before imaging, both plant types underwent a 30-minute dark adaptation period to diminish delayed luminescence. The leaves and plants were then subjected to UPE imaging using the EMCCD camera, with controlled environmental conditions maintained to ensure accurate data acquisition over a 16-hour imaging period for the leaves and 5 minutes imaging for Arabidopsis thaliana, allowing for the observation of UPE variations in response to temperature, injury, and chemical treatments. The Heptapleurum leaves were cut to induce injury at their tips and each leaf was treated with specific chemical agents mentioned above, simulating stress conditions. Arabidopsis thaliana was imaged only at controlled temperatures of 22°C and 26°C for the effect of temperature.

##### 3. Data analysis

After data acquisition from mice and plants, we did not employ any image processing filters to the images. The rest of the analysis was done in the Andor Solis software. However, a color palette was optimized by histogram stretching to achieve maximal contrast of relevant features and image quality. A singular color palette was utilized for visualization of all images in each series of similar experiments.

For the temperature dependence experiments, the region of interest (ROI) was considered to be the entire leaf, while for the injury experiments, two ROIs of equal size were considered, one at the injury site and another in a region on the leaf that was far away from the injury site. The ratio was taken as the quotient of the mean photon counts from these two ROIs on the same leaf and then plotted in Python.

## IV. RESULTS

### 1. UPE from living and dead mice

The death or euthanasia of an animal causes significant physiological changes, including alterations in ROS generation and, hence, should alter the level of UPE. Our experiment involving four mice provides evidence of these changes. Fig.3(A) demonstrate the UPE images from live mice (top row, N=4) and recently euthanized the mice above (bottom row, N=4) following 30 minutes of dark acclimation and a 60-minute imaging session. The UPE imaging results show that the live mice consistently exhibit a significantly higher UPE intensity compared to their deceased counterparts, see Fig. 3(B). This disparity illustrates a profound impact of a system’s vital state on UPE emissions. While the live mice emit robust UPE, likely indicative of ongoing biological processes and cellular activity, the dead mice’s UPE emission is nearly extinguished, with only a few remaining “bright spots”, which correspond to “brighter spots” in living mice, reflecting the cessation of metabolic and energy variation. Note that the body temperatures of both living and deceased mice in the experiments were at 37°C, and the stage supporting the mice was also kept at a temperature of 37°C.

**FIG. 3.**
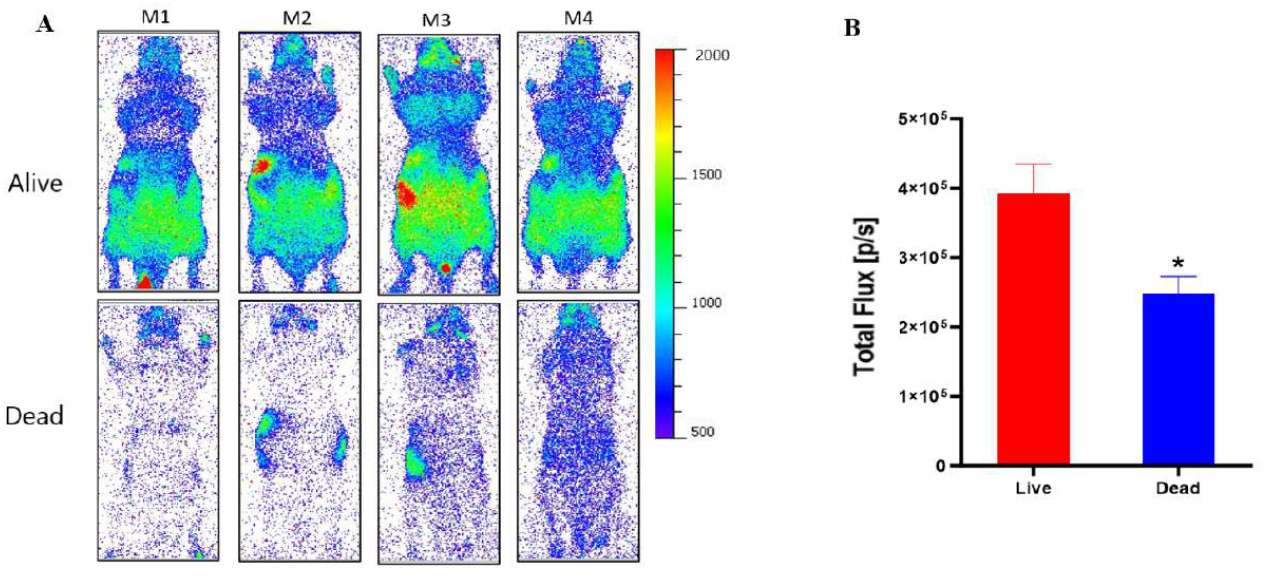
Reduced biophoton emission in euthanized mice compared to live mice. (A) UPE images from live mice (top row, N=4) and recently euthanized mice (bottom row, N=4) following 30 minutes of dark acclimation and a 60-minute biophoton imaging session. (B) Graph showing total photon flux (photons/sec) in live versus recently euthanized mice. Total flux was quantified using a whole-body region of interest (ROI) from the mice in panel A. * Pair t-test, p*<* 0.05. The bodies of both living and deceased mice in the experiments above were at 37°C.

### 2. Effect of temperature on UPE from plant

Temperature plays an important role in modulating biochemical reactions and metabolic processes within cells, including those involved in ROS generation. Generally, higher temperatures tend to accelerate metabolic rates, leading to increased ROS production through processes such as mitochondrial respiration and enzymatic reactions. Additionally, elevated temperatures can disrupt cellular homeostasis, potentially impairing antioxidant defence mechanisms and exacerbating oxidative stress. As for UPE, temperature can influence the rate of molecular excitations and relaxations within cells, consequently affecting photon emission patterns [54]. Higher temperatures may enhance photon emission due to increased metabolic activity and energy transfer processes, while lower temperatures could dampen photon emission rates.

In this experiment, we imaged live Arabidopsis thaliana plants for N=2 replicates and freshly cut green leaves of Heptapleurum arboricola under controlled conditions for N=4 replicates. We conducted the study on Arabidopsis at two different temperatures, 22°C and 26°C, and on cut leaves at six temperatures, from 24°C to 39°C, to observe the variations in UPE intensity. Using EMCCD camera, we captured and analyzed UPE from the plant at these specified temperatures. The results shown in Fig. 4 reveal a significant increase emission intensity at higher temperatures, potentially from enhancement of the biochemical processes that cause UPE. This effect is consistent for all of the observed cut leaves (Fig. 4C). However, a noticeable drop in UPE intensity was observed above 36°C, which could be attributed to the onset of thermal stress, leading to the disruption of cellular integrity and a consequent decline in photon emission.

**FIG. 4.**
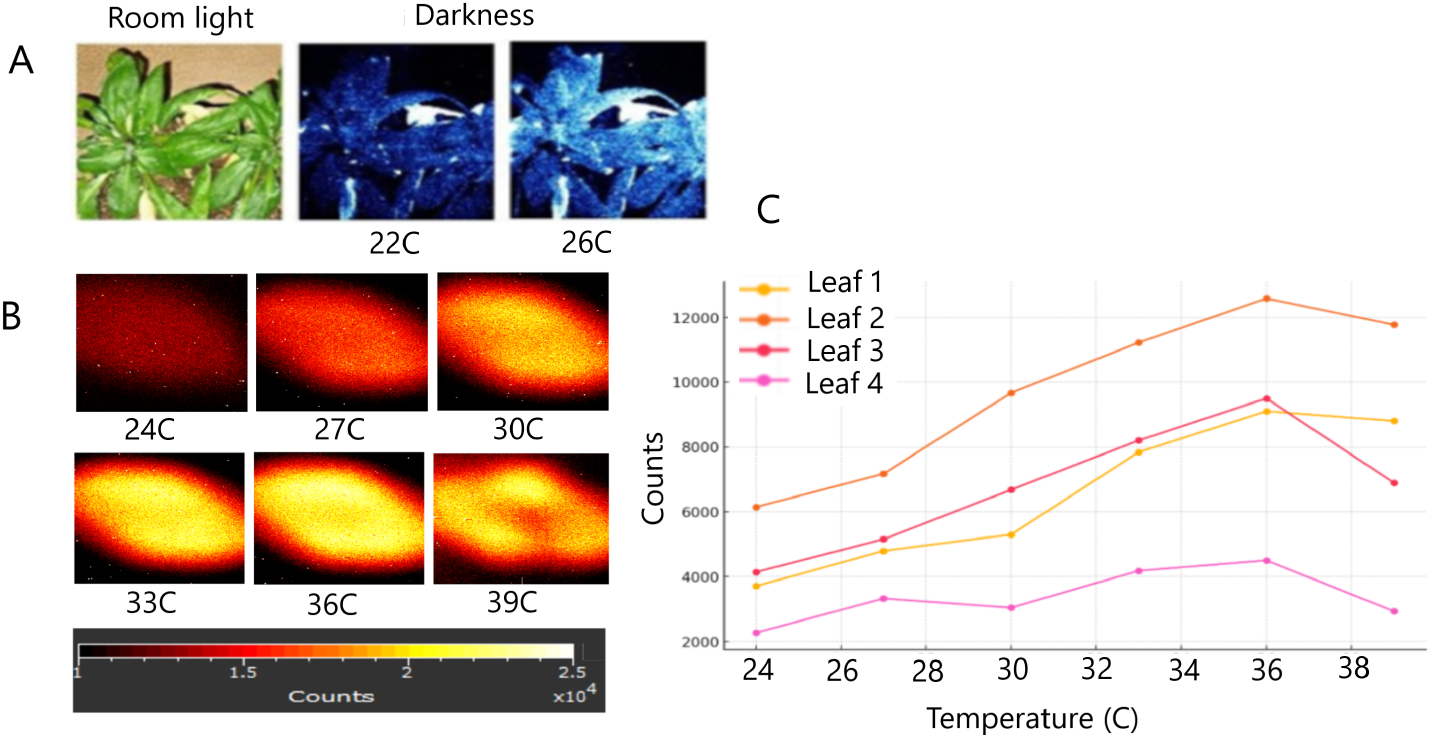
A) Effect of temperature on the intensity of UPE from Arabidopsis thaliana. The exposure time for each image is 5 min. B) Images of UPE from a Heptapleurum arboricola green leaf at different temperatures showing increased intensity as temperatures increase up to 39°C. C) Effect of temperature on UPE from four Heptapleurum arboricola green leaves (N=4) on a temperature-controlled plate where each image had a 15-minute acquisition time. The counts are average of counts over the ROIs over the entire of each leaf. Since the leaf was positioned closer to the lens than the focal length, largely due to being placed on the temperature-controlled plate under the camera, the image resolution is low, so only the intensity variations of UPE are analyzed here. A noticeable drop in UPE intensity was observed after 36°C, probably due to thermal stress disrupting cellular integrity and reducing photon emission [55].

### 3. Injury and chemical effects on leaves

When plants are injured, whether due to physical damage, herbivory, or environmental stress, they activate a series of biochemical and physiological responses. This includes the release of ROS, changes in hormone levels (such as jasmonic acid and ethylene), and altered gene expression to initiate repair and defense mechanisms. These responses are crucial for the plant’s survival and adaptation to stress conditions. This myriad of biological factors may cause a local increase in UPE emission at the injury site [71]. In our study, we observed differential emission intensities for injury sites treated with different chemical agents.

To study the effect of injury, we used Heptapleurum arboricola fresh leaves with cutting effects on the surface (see Fig.5). We also investigated the effects of different chemical agents on the injury response of plants (see Fig. 5). The chemicals that we used were alcohol (Isopropanol), hydrogen peroxide (H_2_O_2_), and benzocaine. In our experiment, four leaves were injured towards their tips. UPE imaging for 16 hours with 1h exposure per image (see Fig.5), with N=5 replicates, each replicate corresponds to a group of four leaves, where three leaves in each group were treated with chemicals, alcohol, hydrogen peroxide, and benzocaine, respectively, and one leaf in each group was left untreated as a regular leaf as control. Our results show that the injury parts in all leaves were significantly brighter than the other parts of the leaves during the 16 hours of imaging. The ratios of photon emission during the first hour from the injury site versus the control were investigated, with each chemical yielding a differential response. The intensity of UPE emission from the injury site treated with benzocaine (with 20% v/v) was far higher than even that of hydrogen peroxide, a well-known enhancer of UPE [43], and higher than alcohol, while benzocaine was dissolved in alcohol. Alcohol is the vehicle and the control of just alcohol justifies that the anesthetic has a different effect on the leaf. The main target of benzocaine in animal cells is Na channels. Plants also have Na channels [39] but there is an uncertainty about the main mechanism of why such an increment in UPE is observed, however, we speculate that the mechanism might be related to ROS [40–42]. Future studies can contribute to an improved understanding of anesthetic action and injury response.

**FIG. 5.**
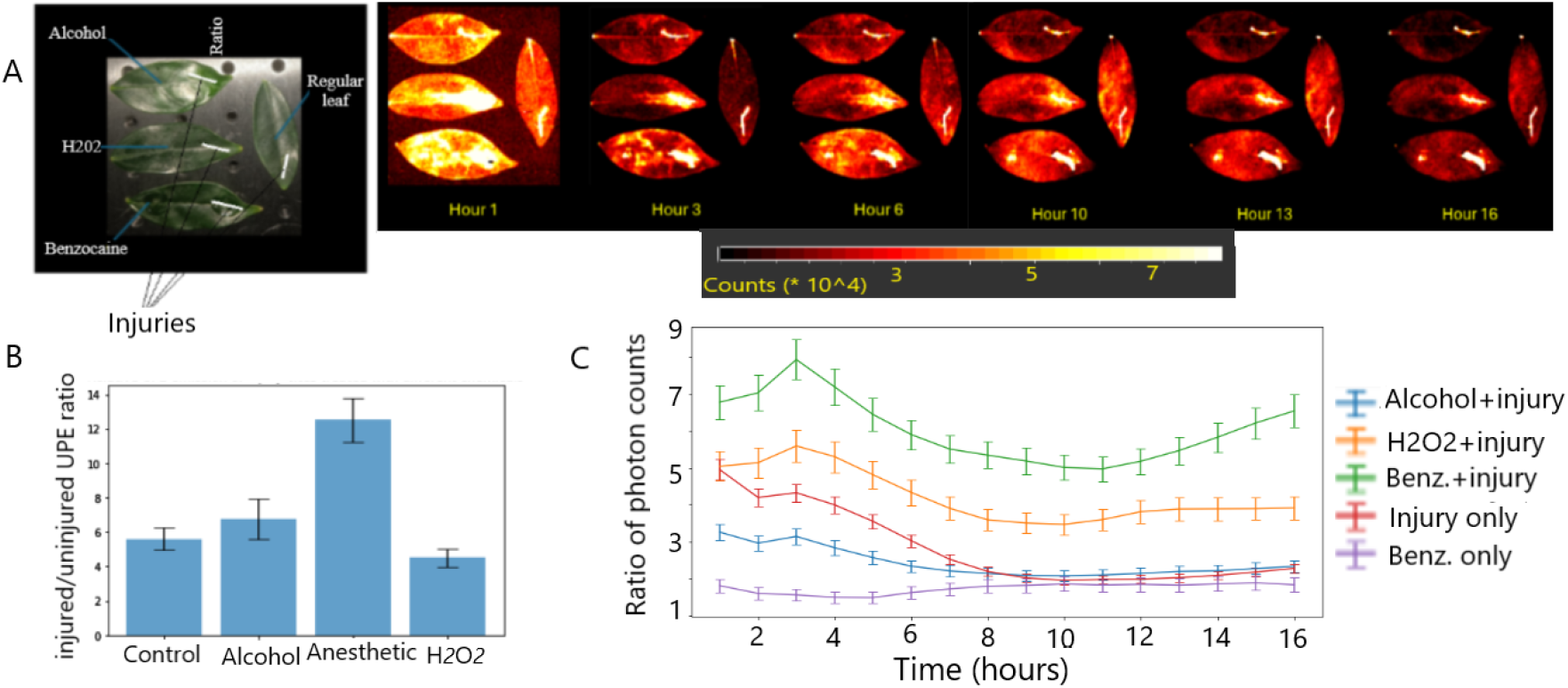
Effect of stress on UPE emissaion from leaves. A) All All Heptapleurum arboricola leaves were injured towards their tip. Images were acquired with 1-hour exposures for 16 hours in total. In each image one leaf at the top left is treated with alcohol, the leaf in the middle is treated with hydrogen peroxide, the leaf at the bottom is treated with benzocaine (anesthetic gel), and the right leaf is not treated with any chemical agent, thus, serving as our control sample. All reagents were applied to the entire surface of the leaf. B) The ratios of intensity between injured and non-injured ROIs in the first hour treated with different chemicals are plotted (Left, bottom), for replicates N=5. C) Time evolution of the variation of the ratios of the injured site relative to the uninjured region in the leaf over 16 hours of imaging. For the comparison of benzocaine independent of injury, it was applied to one side of the leaf while the other side received no treatment. The rest of the chemical treatments were applied on the whole surface of the leaf. (Right, bottom)

To study the effect of injury, we used Heptapleurum arboricola fresh leaves a single controlled cut to the surface (see Fig.5). We imaged samples of four leaves for 16 hours with 1-hour exposure per image (see Fig.5), with N=5 replicates, each corresponding to a group of four leaves. Of these, three leaves in each group were treated with chemicals, alcohol (Isopropanol), hydrogen peroxide (H_2_O_2_), and benzocaine, respectively, and one leaf in each group was left untreated as a control. Our results show that the injury parts in all leaves were significantly brighter than the uninjured parts of the leaves during all 16 hours of imaging. The ratios of photon emission during the first hour from the injury site ROI versus the control ROI were investigated, with each chemical yielding a differential response. The intensity of UPE emission from the injury site treated with benzocaine was far higher than even that of hydrogen peroxide, a well-known enhancer of UPE [43], and higher than alcohol. This suggests that the anesthetic could potentially interact with ROS accumulating at the injury site. Future studies can contribute to an improved understanding of anesthetic action and injury response.

## V. SUMMARY AND CONCLUSION

Our study looked at what faint light emissions, called ultraweak photon emission (UPE), can tell us about mice’s vitality and plants’ health under different stressors. Using advanced EMCCD and CCD cameras and imaging mice and plants under the abovementioned conditions, we observed that the bodies of live mice exhibit a greater level of photon emission than deceased mice. This shows in a very clear way that UPE is associated with being alive. Also, the intensity of UPE from plants is influenced by temperature, where higher temperature causes higher UPE while it displays distinct characteristics from blackbody radiation. Moreover, UPE may be used as an indicator of injury effects in plants, where the injured parts emit more photons. The intensity of UPE is also affected by chemical treatments. The effect of anesthetic on injury shows the highest emission compared to others, which needs further investigation to elucidate the mechanism at the cellular level.

The study of UPE has fundamental importance for basic research, as it can be used for non-invasive insights into the biochemical and metabolic processes of living organisms. UPE is closely linked to vitality, with living systems emitting higher levels of photons. Moreover, the influence of environmental factors can affect UPE and therefore it can serve as a valuable indicator of oxidative stress and health abnormalities. This makes UPE a promising tool for advancing our understanding of both plant biology and biomedical research at the fundamental level.

For applications, the investigation of UPE from plants may be used as a simple method for non-invasive monitoring of health abnormality and plant growth under different environmental conditions for plant biology and agricultural practices. The study of UPE from mice bodies can be useful for biomedical research and clinical applications [58–65, 67], particularly for non-invasive label-free imaging of biochemical activities and metabolism in living systems.

## VI. DATA AVAILABILITY STATEMENT

Data are available based on request.

## ACKNOWLEDGMENTS

We acknowledge support from the National Research Council of Canada through its Quantum Sensors Challenge Program, from the Natural Sciences and Engineering Research Council through its Discovery Grant Program and the Alliance quantum consortia grant “Quantum Enhanced Sensing and Imaging”, and from the New Frontiers in Research Fund. We are thankful for fruitful discussions with Hadi Zadeh-Haghighi, Rishabh, Gabriel Bertolesi, and Douglas Muench.

## VIII. CORRESPONDENCE

